# Different lanthanide elements induce strong gene expression changes in a lanthanide-accumulating methylotroph

**DOI:** 10.1101/2023.03.06.530795

**Authors:** Linda Gorniak, Julia Bechwar, Martin Westermann, Frank Steiniger, Carl-Eric Wegner

## Abstract

Lanthanides are the most recently described life metals and are central to methylotrophy in diverse taxa. We recently characterized a novel, lanthanide-dependent, and lanthanide-accumulating methylotroph, Beijerinckiaceae bacterium RH AL1, that utilizes lighter lanthanides (La, Ce, Nd) for methanol oxidation. We show that lanthanum concentration and different lanthanide (Ln) elements strongly affect gene expression and intracellular lanthanide accumulation. Differential gene expression analysis based on incubations with either La (50 nM or 1 µM), Nd (1 µM), or a lanthanide cocktail ([La, Ce, Nd, Dy, Ho, Er, Yb], equimolarly pooled, 1 µM), revealed that up to 41% of the encoded genes were differentially expressed. The effects of lanthanum concentration and Ln elements were not limited to lanthanide-dependent methanol oxidation but reached into many aspects of metabolism. We observed that lanthanides control the flagellar and chemotactic machinery and that they affect polyhydroxyalkanoate (PHA) biosynthesis. Secretion and various uptake systems, and carbohydrate metabolism were highly responsive. The most differentially expressed genes encode various unknown or hypothetical proteins, but also *lanM*, coding for the well-characterized lanthanide-binding protein lanmodulin, and a glucose dehydrogenase gene linked to the conversion of β-D-glucose to gluconolactone, a known metal chelator. Electron microscopy, together with RNAseq, suggested different and potentially selective mechanisms for the uptake and accumulation of individual Ln elements. Mechanisms for discriminating lanthanides and links between lanthanides and various aspects of metabolism underline a broader functional role for lanthanides, possibly by functioning as calcium complements or antagonists.

**Importance:** Since its discovery, lanthanide-dependent metabolism in bacteria attracted a lot of attention due to its bio-metallurgical application potential regarding lanthanide recycling and circular economy. The physiological role of lanthanides is mostly studied dependent on presence and absence. Comparisons of how different (utilizable) lanthanides affect metabolism have rarely been done. Our research shows that strain RH AL1 distinguishes different lanthanide elements and that the effect of lanthanides reaches into many aspects of physiology, for instance, motility and polyhydroxyalkanoate metabolism. Numerous differentially expressed genes coding for unknown or hypothetical proteins might hide so far unknown lanthanide-binding proteins. Our findings regarding lanthanide accumulation suggest different mechanisms for dealing with individual lanthanide elements and provide insights relating to intracellular lanthanide homeostasis. Understanding comprehensively how microbes distinguish and handle different lanthanide elements is key for turning knowledge into application regarding lanthanide-centered biometallurgy.

## INTRODUCTION

Lanthanides (Ln) have been dubbed vitamins of the 21^st^ century due to their relevance for high-tech applications that define our modern, everyday life (1). In nature, lanthanides primarily occur in poorly soluble phosphate and carbonate minerals (2–5), making accessing and recovering them challenging. Lanthanides are also “life metals” (6–8) with high relevance for carbon cycling and methylotrophy. Ln-dependent metabolism is centered around pyrroloquinoline quinone (PQQ)-dependent alcohol dehydrogenases (ADHs). The catalytic activity of PQQ ADHs is based on a metal:PQQ cofactor complex. PQQ ADHs are diverse enzymes, including, among others, Mxa-type methanol dehydrogenases (MDHs) and five clades of Xox-type MDHs (9). Mxa-type, calcium-dependent MDHs were previously considered essential for microbes utilizing C_1_-substrates such as methane or methanol (10–12). Xox-type MDH from *Methylorubrum extorquens* AM1 was the first identified and characterized Ln-dependent enzyme (13).

Genes encoding Xox-type MDH are widely distributed in the environment (14–17), suggesting that methylovory (the supplemental use of C_1_ compounds as energy sources) is more common than previously thought (18–20). ExaF from *M. extorquens* AM1 was the first known Ln-dependent PQQ ADH acting on multicarbon substrates (21). Related enzymes have been identified in the non-methylotroph *Pseudomonas putida* KT2440 (22) and the facultative methylotroph Beijerinckiaceae bacterium RH AL1 (23). Characteristic amino acid residues involved in lanthanide coordination indicate that most PQQ ADHs are lanthanide-dependent (9,24). The unknown substrate spectrum of many enzymes suggests a broader role for lanthanide-dependent metabolism.

Ln-utilizing microbes must be able to access, mobilize, and take up lanthanides, despite potentially low bioavailability. Methylotrophs studied have a preference for lighter lanthanides (La-Nd). Heavier lanthanides are generally less favored or not utilized (6,7). For *M. extorquens,* it was shown that lanthanide uptake is enabled by a transport system comprising a TonB-dependent receptor (LutH) and an ABC transporter (LutAEF), which are encoded in the *lut*-cluster (lanthanide <u>u</u>tilization and transport) (25). LutH is responsible for periplasmic uptake, while LutAEF facilitates cytoplasmic uptake. The first identified and best-studied lanthanide-binding protein, besides PQQ ADHs, was lanmodulin (LanM). LanM is a homolog of the well-characterized calcium-binding protein calmodulin (26), which features high affinities for lanthanides and application potential for lanthanide detection and recovery (27–29). Few other lanthanide-binding proteins have been identified, including most recently lanpepsy from *Methylobacillus flagellatus* (27,30,31). Intracellular accumulation of lanthanides was shown for *M. extorquens* AM1 *(25)* and Beijerinckiaceae bacterium RH AL1 (32). *M. extorquens* stores lanthanides in the cytoplasm, while strain AL1 keeps periplasmic lanthanide deposits. An evolved *M. extorquens* strain that hyperaccumulates gadolinium was recently described (33). RNAseq analyses of *M. extorquens* grown with soluble and less soluble Ln led to the identification of a gene cluster linked to lanthanophore biosynthesis (34).

We used Beijerinckiaceae bacterium RH AL1 (23,32) to study lanthanide discrimination and accumulation. Up to 41% of the encoded genes were differentially expressed under methylotrophic growth conditions when La was swapped for Nd, or an equimolarly pooled cocktail of light and heavy lanthanides (La, Ce, Nd, Dy, Ho, Er, Yb). Electron microscopy showed that strain AL1 accumulates Nd, as shown before for La (32), in the periplasm. Periplasmic storage was also visible for the lanthanide cocktail. Ln elements were differently accumulated, supporting the idea of selective lanthanide uptake (32). We could show that the effects of lanthanum concentration and different lanthanide elements stretched way beyond the lanthanome. We hypothesize that lanthanides interfere with or complement the physiological role of calcium.

## RESULTS

### Lanthanum concentration, lanthanide elements, and their effects on growth

We cultivated Beijerinckiaceae bacterium RH AL1 with different lanthanum concentrations (50 nM vs. 1 µM) or 1 µM of different lanthanide elements (La, Nd, or an Ln cocktail [La, Ce, Nd, Dy, Ho, Er, Yb; equimolarly pooled]) using 0.5% methanol (v/v) as the carbon source. We noted significantly (p ≤ 0.05, *Student’s t-test*) different growth rates for 50 nM and 1µM cultures (0.038 ± 2.13×10 ^-4^ h^-1^ and 0.044 ± 0.001 h^-1^) (**Figure 1A**, left panel). Cell numbers increased from 2.22 ± 3.79 ×10^6^ and 2.47 ± 3.49 × 10^6^ ml^-1^ (t_0_) to 1.49 ± 0.71 × 10^9^ and 2.64 ± 0.51 × 10^9^ ml^-1^ (t_2_) (**Figure 1B**, left panel; **Table S1**). The incubations with different lanthanide elements revealed overall comparable growth patterns (**Figure 1A**, right panel), but cultures grown with lanthanum showed, compared to neodymium cultures, significantly (p ≤ 0.05) faster growth with a doubling time of 16.70 ± 0.46 h and a growth rate of 0.042 (h^-1^). Cell numbers increased up to 2.41 - 3.68 × 10^9^ ml^-1^ (t_2_) for the different setups (**Figure 1B**, right panel; **Table S1**).

**FIG 1.**
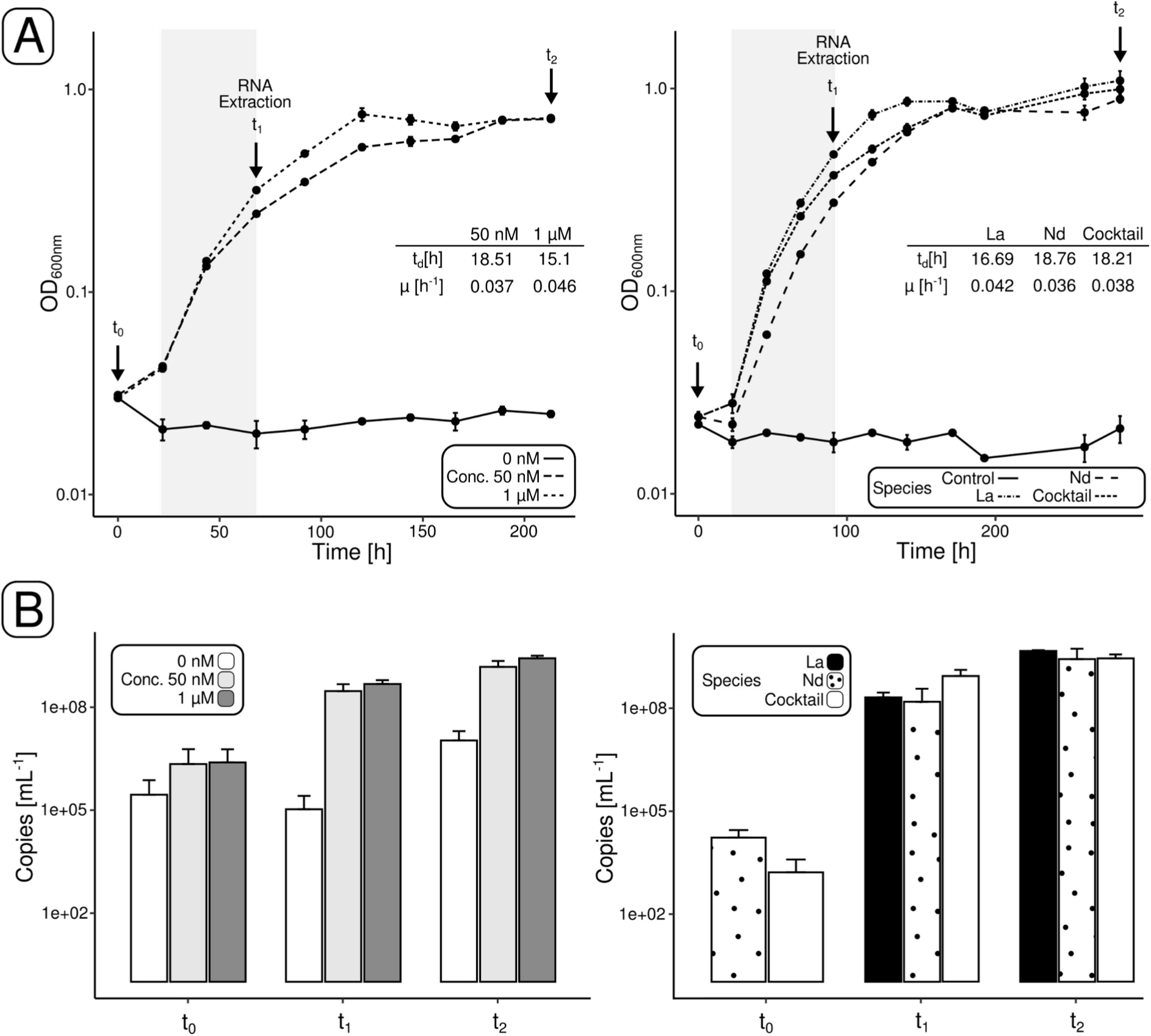
Growth of Beijerinckiaceae bacterium RH AL1 with different concentrations of lanthanum and different lanthanide elements. Growth was monitored spectrophotometrically (A) and by 16S rRNA gene-targeting quantitative PCR. The grey shade in (A) indicates the time interval considered for calculating doubling time (t_d_) and growth rate (µ). t_0_-t_2_ = time points for qPCR and RNA extraction (t_1_).

### Overall transcriptome changes

Biomass from the mid-exponential phase (t_1_, **Figure 1A**) was used for RNAseq (**Table S2**). Genes with changes in the expression above |0.58| log_2_FC (fold change) and expression values higher than 4 log_2_CPM (counts per million) (**Figures S1-4**) were considered for DGEA (differential gene expression analysis) (**Figure 2A**, **Table S3-6**). We made the following comparisons: (1) 50 nM La vs. 1µM, (2) 1µM La vs. 1µM Nd, (3) 1µM La vs. 1µM Ln cocktail, and (4) 1µM Nd vs. 1µM Ln cocktail (**Figure 2A**). Up to 41% of the genome was differentially expressed in the case of (2) and (3) (**Figure 2B**). Gene expression differed less for (4) and (1). We identified 320 and 351 DEGs (differentially expressed genes), representing 7.4 and 8.12% of the genome, respectively.

**FIG 2.**
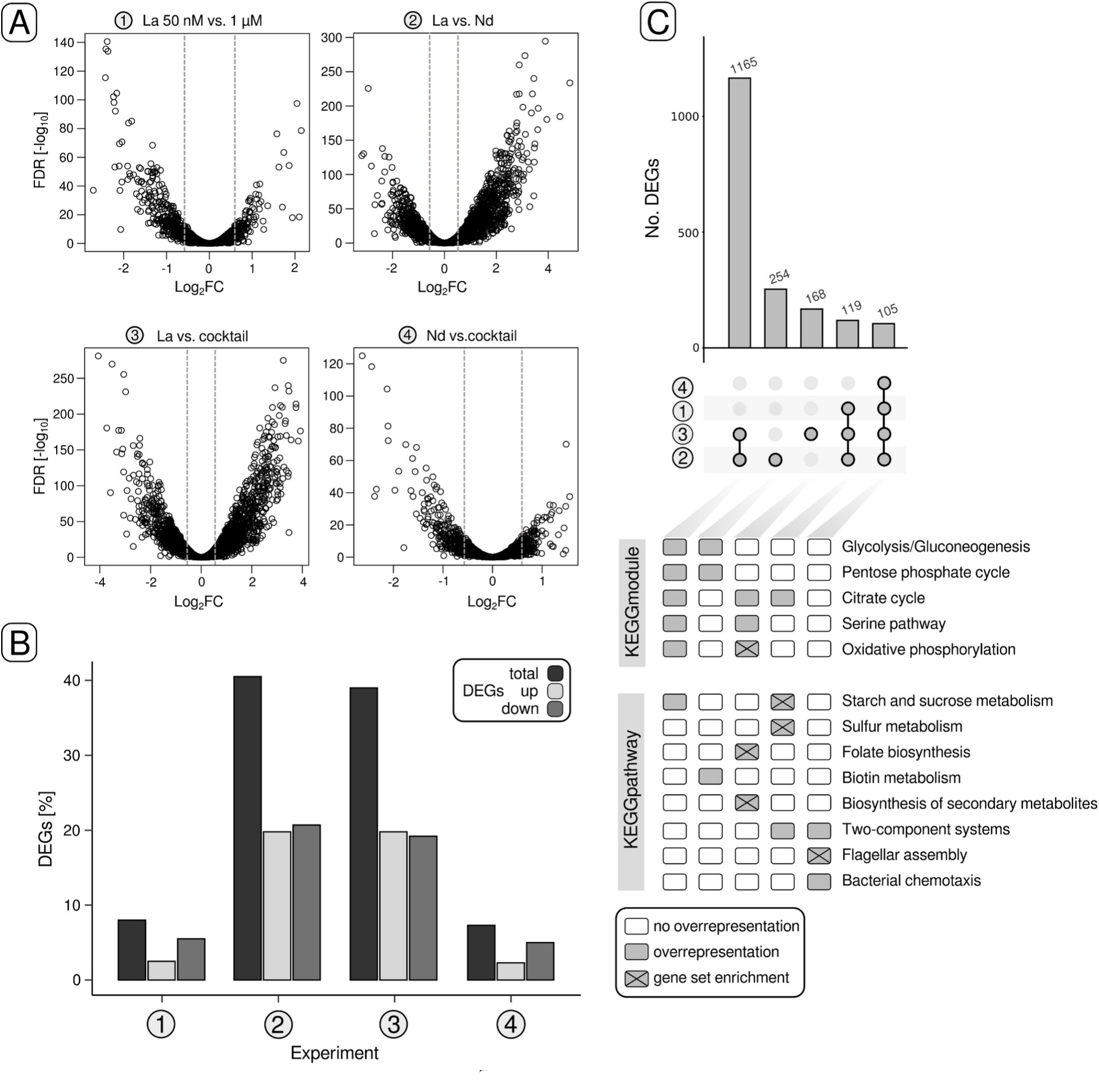
Differential gene expression in response to lanthanum concentration and lanthanide elements. Volcano plots indicate differentially expressed genes (DEG) for the four different comparisons (1)-(4) (A). Genes with changes in gene expression above |0.58| log_2_FC (fold change), gene expression values higher than 4 (log_2_CPM, counts per million) and p-values smaller than 0.05 were considered for downstream analysis. The grey dashed lines indicate the Log_2_FC threshold of |0.58|. The proportions of total DEGs, up-and downregulated genes in relation to the number of genes encoded in the genome are shown as a bar chart (B). UpSet plots were used to highlight intersections, meaning shared genes, between the different comparisons (C). The lower half of (C) indicates potential overrepresentation and enrichment of KEGGmodules and KEGGpathways. FDR = false-discovery rate.

Taking a closer look at the DEGs in AL1 revealed a substantial overlap of 1165 genes between (2) and (3) (**Figure 2C**). Independent of the comparison, a subset of 105 genes was differentially expressed in all cases. Overrepresentation and gene set enrichment analysis based on KEGG annotations revealed an overrepresentation of genes linked to flagellar assembly, two-component systems, and bacterial chemotaxis (**Figure 2D**, **Table S7**). KEGG modules related to central carbohydrate and energy metabolism were overrepresented in the 1165 overlapping genes between (2) and (3).

### Differentially expressed genes and pathways

Motility-and chemotaxis-related genes were downregulated in all comparisons (**Figure 3**, **Table S8**). In the case of (2) and (3), log_2_FC values for flagellar genes ranged from −0.60 to −4.27 (log_2_CPM values between 4.25 and 9.33). We investigated the effect of lanthanide concentration (5 nm −10 µM) and elements (La, Nd, Ln cocktail) on motility by employing a soft agar and TTC (2,3,5−triphenyltetrazolium chloride)-based assay (**Figure S5**, **Table S9**). We observed a decrease in motility with increasing lanthanide concentration. Motility was higher for cultures grown with Nd or the Ln cocktail than La.

**FIG 3.**
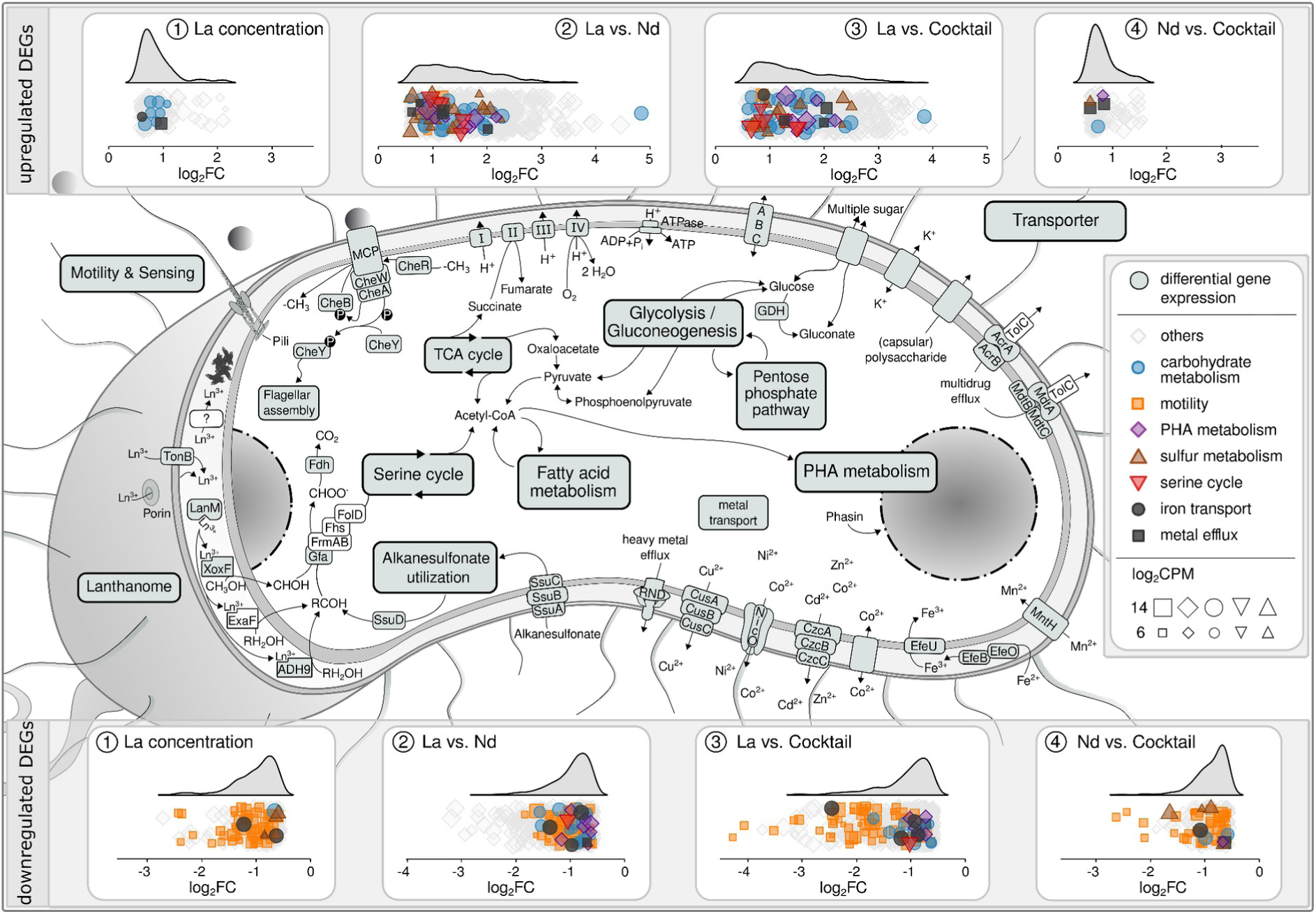
Graphical representation of differential gene expression. Aspects of metabolism that responded (in part) to lanthanide elements and/or lanthanum concentration are summarized (based on **Table S8**). For each comparison, up-and downregulation of genes are illustrated through ridge plots based on log _2_FC values. Marker sizes correspond to the log_2_CPM values. Selected genes associated with motility, carbohydrate metabolism, PHA metabolism, serine cycle, sulfur metabolism, iron transport, and metal efflux were highlighted.

**FIG 4.**
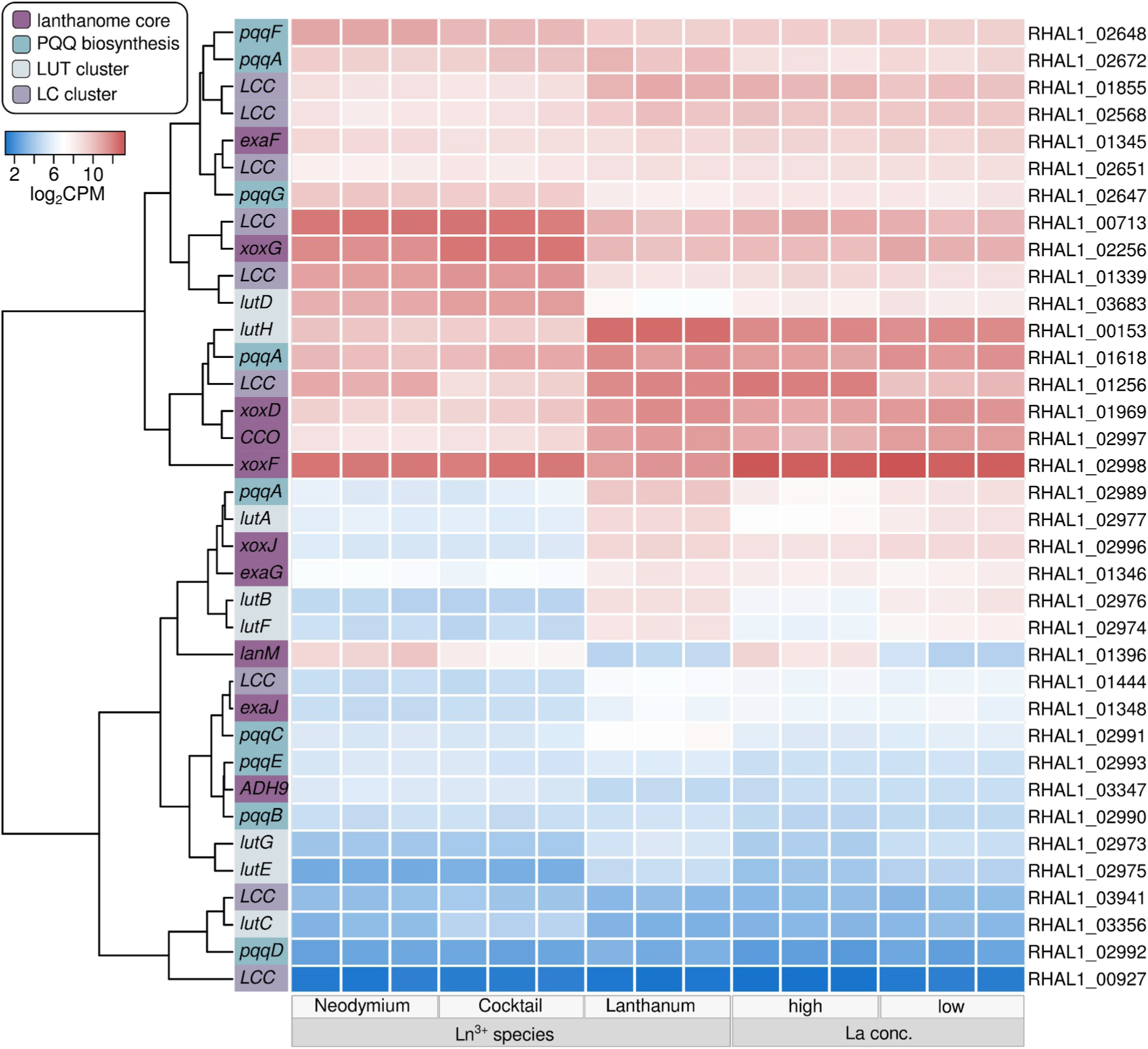
Gene expression of lanthanome (-related) genes. We defined four groups of genes linked to the lanthanome: lanthanome core, PQQ biosynthesis, and *Lut*−cluster The core lanthanome comprises genes for key enzymes involved in lanthanide-dependent methanol oxidation. Lut and LC clusters refer to identified gene homologs of the previously described *Lut*-cluster (lanthanide utilization and transport) lanthanide chelator cluster (25,34). The row dendrogram is based on euclidean distances. The colored vertical sidebar shows to which of the four outlined groups the respective gene belongs. Gene expression is given in log _2_CPM (counts per million) and indicated by a blue-red color scale.

Genes encoding proteins (SsuABCD) enabling alkanesulfonate uptake and utilization were downregulated when strain AL1 was grown with 1 µM (log_2_FC −0.59 and −0.84, log_2_CPM 4.25 - 7.40) instead of 50 nM La, but upregulated in Nd and Ln cocktail samples (**Table S4+5**). Assimilatory sulfate reduction genes were upregulated with respect to Nd. We noticed numerous genes associated with heavy metal efflux, iron transport, and polysaccharide export, as well as genes for porins (**Table S8**). The latter were downregulated upon increased La concentration and when La was swapped for Nd or the Ln cocktail. Genes coding for the iron-storage protein bacterioferritin (RHAL1_00987) and EfeUO (RHAL1_03444, RHAL1_03445), part of the EfeUOB iron uptake system, were downregulated in the case of (2) and (3). The same was true for the *ebbB* and *ebbD,* genes that encode the energy transmission machinery for TonB-dependent uptake.

A gene (RHAL1_01212) encoding a glucose-1 dehydrogenase was the (second) most upregulated (log_2_FC 4.83 and 3.84, log_2_CPM 10.08) in the case of (2) and (3). Genes coding for hypothetical and unknown proteins made up more than 50% of the top 100 upregulated genes when strain AL1 was grown with the Ln cocktail instead of lanthanum. The most upregulated gene (RHAL1_03682) coding for a hypothetical protein yielded no hits when searching it against NCBI nr, and Pfam. Identifying protein domains with Interpro suggested that the encoded protein is cytoplasmic and partially membrane-embedded. Single genes of central carbohydrate metabolism pathways were differentially expressed and predominantly upregulated (**Figure 3**, **Table S8**) in response to Nd and the Ln cocktail, including succinate dehydrogenase (flavoprotein subunit), malate dehydrogenase, and fumarate hydratase genes. Formate dehydrogenase genes were slightly upregulated with increased lanthanum concentration (RHAL1_03901-03903, log_2_FC 0.72-0.88). Gene expression linked to C_1_-assimilation through the serine cycle was affected by swapping the lanthanide element. The gene encoding serine hydroxymethyltransferase (RHAL1_01794) was upregulated when strain AL1 was grown with either Nd or the Ln cocktail, *sga* (serine-glyoxylate aminotransferase, RHAL1_01353) was higher expressed when AL1 was supplemented with Nd. At the same time, *hpr* (hydroxypyruvate reductase, RHAL1_03822) was less expressed.

Cultivation with Nd and the Ln cocktail caused differential expression of PHA (polyhydroxyalkanoate)-(de)polymerization-related genes. A gene encoding the PHA synthesis repressor PhaR (RHAL1_03606) was downregulated in Ln and Nd samples (log_2_FC −1.16 and −1.02). Two genes (RHAL1_01143 and RHAL1_3443), coding for phasins (proteins with a scaffolding role in PHA granule formation), were highly expressed and upregulated in Nd and Ln cocktail cultures (log_2_FC 0.90 and 1.76, log_2_CPM 12.22 and 13.62). Strain AL1 carries four polyhydroxyalkanoate depolymerase genes. Two were up− (RHAL1_03589, RHAL1_01066), and two downregulated (RHAL1_01171, RHAL1_01370) with log_2_FC values ranging from − 0.68 to 2.19 (log_2_CPM between 4.97 and 8.65).

### The lanthanome in response to lanthanide elements and concentration

Beijerinckiaceae bacterium RH AL1 encodes three PQQ ADHs, one clade 5 XoxF MDH, ExaF ADH, and a subgroup 9 PQQ ADH. The *xoxF5* (RHAL1_02298) gene was moderately upregulated in response to Nd and the Ln cocktail (log_2_FC 1.13 and 1.04). Gene expression was high across all conditions (**Table S10**). The same held true for the gene encoding XoxG (RHAL1_02256), the complementary cytochrome c_L_ of XoxF. The expression of *exaF* (RHAL1_01345) did not differ significantly when comparing conditions. RHAL1_03347, encoding the subgroup 9 PQQ ADH, was slightly upregulated in response to Nd (0.74). Differences in lanthanum concentration led to a strong downregulation (log_2_FC −2.71) of *lanM* (RHAL1_01396), while Nd and the Ln cocktail triggered upregulation (log_2_FC 3.29, 2.08). Comparing Nd and Ln cocktail incubations showed downregulation of *lanM* (log_2_FC −1.20).

Most of the gene homologs (**Table S11**) of the *lut*-cluster and lanthanide chelator (LC) cluster were downregulated when comparing Nd and the Ln cocktail to La. This was most obvious for *lutH*, which encodes the TonB-dependent receptor involved in periplasmic lanthanide uptake (log_2_FC −3.11, −3.34). The gene homologs (RHAL1_02974, RHAL1_02975, RHAL1_02977) of the ABC transporter LutAEF, crucial for cytoplasmic lanthanide uptake, were upregulated when strain AL1 was grown with 1 µM La in comparison to 50 nM. The gene (RHAL1_3683) coding for the periplasmic lanthanide-binding protein LutD was upregulated in Nd and Ln cocktail samples. RHAL1_01256, the gene homolog (**Table S3-6**) of the TonB-dependent receptor of the LC cluster, was downregulated across all comparisons.

The three copies of *pqqA* (RHAL1_01618, RHAL1_02672, RHAL1_02989) were downregulated in response to Nd and the Ln cocktail. The degree of downregulation differed (log_2_FC [RHAL_01618] −1.43, −1.91; [RHAL1_02672] −0.68 [RHAL1_02989 −2.68, −2.60). Likewise, *pqqC* was also downregulated in Nd and Ln cocktail incubations, while *pqqFG* (RHAL1_02647, RHAL1_02648) were moderately upregulated.

### Differences in intracellular lanthanide deposition

Screening ultrathin sections from cultures grown with Nd or the lanthanide cocktail revealed peripheral, periplasmic deposits in the proximity of the cell poles and close to polyhydroxyalkanoate (PHA) vacuoles (**Figure 5A**, upper left + upper middle panel). FFTEM confirmed the localization of these intracellular deposits (**Figure 5A**, upper right panel). We revisited ultrathin sections from previous work (32) and compared them to ultrathin sections from cultures grown with Nd and the Ln cocktail to analyze the effect of swapping lanthanide elements on PHA biosynthesis (**Figure 5B**). The average cell area when strain AL1 was grown with La was 1.09 ± 0.31 µm^2^, while the respective values were significantly (*p* ≤ 0.001, *t-test*) higher for Nd (1.79 ± 0.58 µm^2^, +64%) and the Ln cocktail (1.74 ± 0.53 µm^2^, +59%), respectively (**Figure 5B**). PHA vacuoles occupied between 0.11 ± 0.04 (La) and 0.58 ± 0.20 (Nd) µm ^2^, which was equivalent to between 10.5 ± 4.9 and 32.9 ± 5.2% of the cell area. The areas occupied by PHA vacuoles were 3.13− and 2.43 times bigger for Nd and the Ln cocktail than for La (**Figure 5B**).

**FIG 5.**
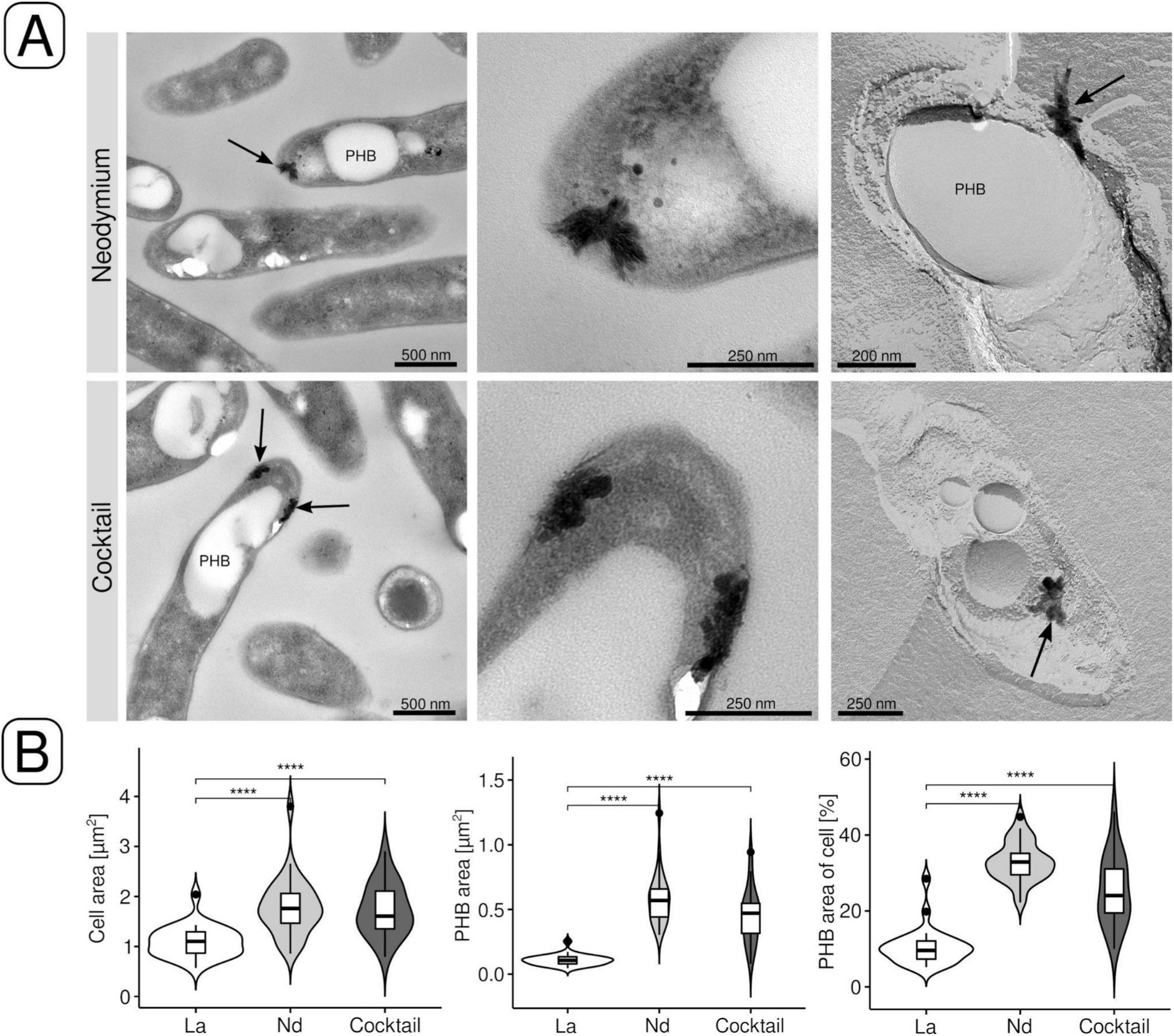
Electron microscopic examination of periplasmic lanthanide deposits. Deposits were identified by transmission electron microscopy (TEM) (A) (upper left panels). The upper middle panels are close-ups of the areas shown in the upper left panels. Periplasmic deposits were also identified by freeze-fracture TEM (FFTEM) (upper right panels). The sizes of cells grown with La (32), Nd, or the Ln cocktail were compared by measuring cell areas. We also compared the area occupied by polyhydroxybutyrate (PHB) vacuoles based on the area and the area of the cell occupied by them (B). The analysis was done for three images (magnification: 4000 ✕, image area: 540.5 µm^2^), and between 27 to 30 cells were analyzed per condition (La, Nd, cocktail).

We verified that the identified periplasmic deposits contained lanthanides through energy-dispersive X-ray spectroscopy (EDX) starting from ultrathin sections and freeze-fracture replicas (**Figure 6A+B**). Concerning the samples originating from Ln cocktail incubations (**Figure 6B**), distinct signals were detected for Ce, Nd, Dy, Ho, and Er, but not for La and only weakly for Yb. The share of the different lanthanides in the deposits (**Table S10**) ranged between 1.6% (ytterbium) and 30.4% (cerium). The ratio between P and Ln was between 0 (n = 3, EM data from (32)) for La deposits and 12 for Nd and Ln deposits (n = 3) (**Figure S6**, **Table S12**).

**FIG 6.**
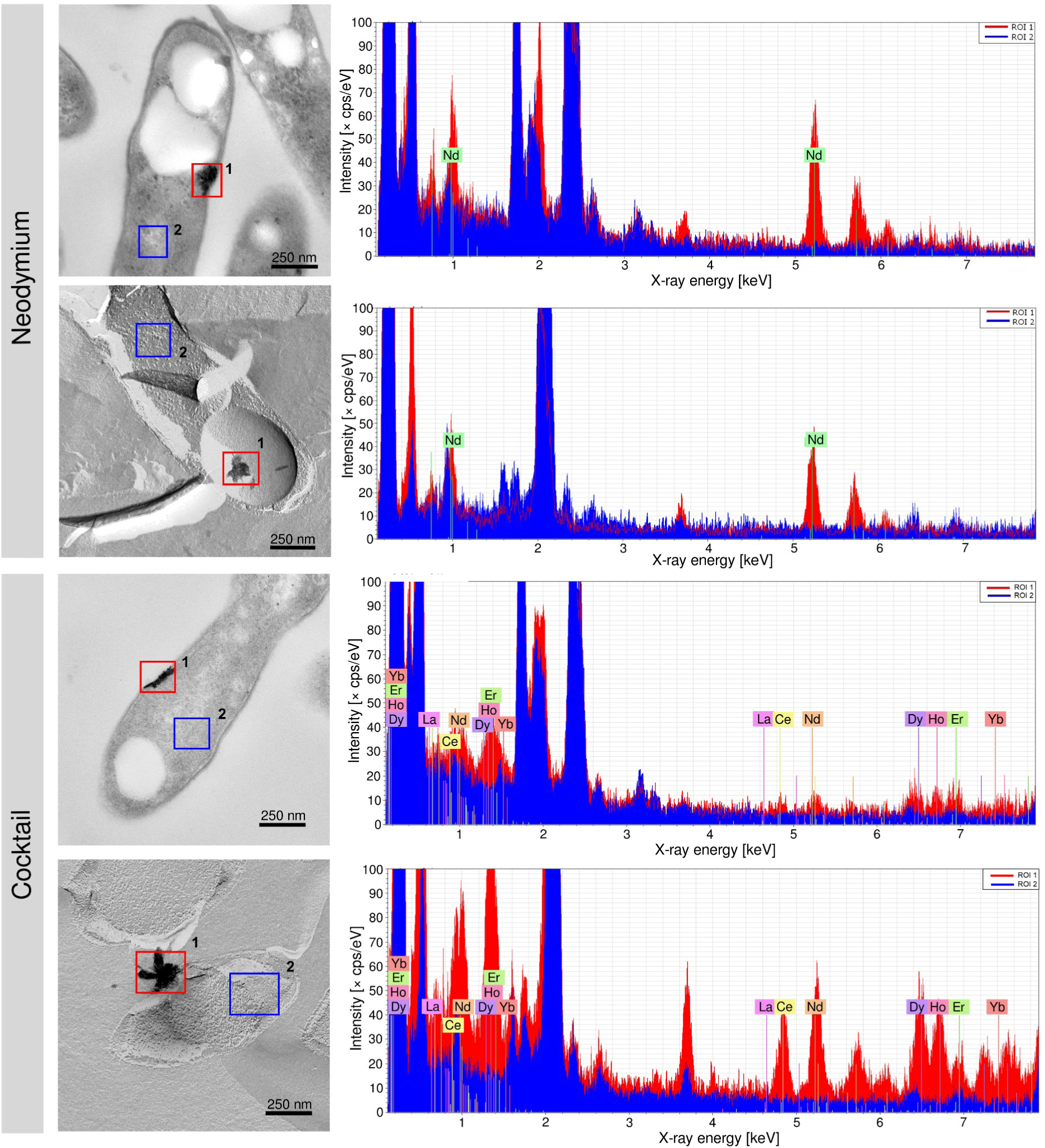
Elemental analysis of periplasmic lanthanide deposits. TEM and FFTEM specimens were subjected to elemental analysis based on energy-dispersive X-ray spectroscopy (EDX). Red (deposit) and blue (reference) boxes indicate measured areas ROI = region of interest, cps = counts per second. Scale bar = 250 nm.

## DISCUSSION

Minor differences in growth masked pronounced gene expression changes caused by different lanthanide elements. Beijerinckiaceae bacterium RH AL1 is apparently able to sense and distinguish different (light and utilizable) lanthanide elements. The effect of lanthanides extended beyond the lanthanome and C_1_-metabolism, indicating a broader role for lanthanides. Past studies reported positive effects on growth, dependent on the presence or absence of lanthanides in (non-)methanotrophic methylotrophs that possess calcium-and lanthanide-dependent MDH (35–40). Similar to our findings here, only small differences were reported when comparing the effect of different light lanthanide elements on growth (38,41).

Few studies previously addressed gene/protein expression changes in response to lanthanide supplementation (36,38–40) and only in microorganisms featuring Mxa-and Xox-type MDHs. It seemed as if lanthanides control small numbers of genes, mostly *mxa*−and *xox*-cluster genes, in organisms that encode the lanthanide switch. *Methylobacterium aquaticum* 22A is the only organism for which the effect of different lanthanides (La, Ho, Lu) on gene expression was tested. Only La affected the gene expression of methylotrophy-related genes. Ho and Lu did not trigger the lanthanide switch, likely because of the preference for light lanthanides (38).

Observed differences between the La, Nd, and Ln cocktail treatments support that aspects of physiology affected by lanthanides are tuned towards certain lanthanide elements. In *M. buryatense* 5GB1C, lanthanum and cerium triggered the lanthanide switch, but lanthanum had a stronger effect on *xoxF* expression (42). Thermal stability analysis showed that XoxF metallation in *M. extorquens* AM1 affects the integrity and that La is preferred over Nd (43). The catalytic efficiency of XoxF was affected by metallation in *Methylacidiphium fumariolicum* SolV and higher for lighter lanthanides (44). The observed upregulation of *xoxF* and *xoxG* in the case of strain AL1 could have compensated for a potentially reduced catalytic efficiency with Nd. Differences in ionic radius, Lewis acidity, and redox potential likely determine the widespread preference for lighter lanthanides. The latter affects the physiological electron acceptor of XoxF, XoxG, which complements XoxF with a reduction potential tailored towards lanthanide elements that are ideally suited for XoxF (45). It is known from iron homeostasis (46,47) that TonB-ABC transport systems are commonly downregulated if iron is sufficiently available. The same was shown for *lutH* and *lutAEF,* which encode a TonB-ABC transport system for lanthanide uptake into peri-and cytoplasm (25,48). We noted the same pattern in this work and also previously in a non-methylotrophic background (32). The downregulation of PQQ biosynthesis genes when strain AL1 was grown with Nd and the Ln cocktail, especially *pqqA* copies, seems counterintuitive. XoxF requires PQQ as a cofactor and *xoxF* was upregulated. Similar observations were made in *M. aquaticum* 22A when comparing methylotrophic growth with Ca and La (38).

Lanmodulin was the first known lanthanide-binding protein (except PQQ ADHs) (26). We meanwhile know three more: the periplasmic Ln-binding protein LutD; supposed to be associated with the LutAEF ABC transporter (27), a ubiquitin ligase from plant chloroplasts (30), and most recently lanpepsy (31) from *Methylobacillus flagellatus*. Lanmodulin is not essential for lanthanide-dependent methylotrophy in *M. extorquens* (25). The responsiveness of *lanM* that we observed is not restricted to methylotrophic growth. We previously showed that *lanM* was the most differentially expressed gene in strain AL1 when comparing growth with pyruvate as a carbon source in the presence and absence of La (32). Like calmodulin and calcium, lanmodulin might play a key role with respect to lanthanide homeostasis.

Calcium affects many aspects of bacterial physiology. Our findings suggest that lanthanides might do so too - as calcium analogs/mimics or antagonists. The role of calcium as a regulator and secondary messenger is well-established in eukaryotes (49–51), but poorly understood in prokaryotes (51,52). Multiple aspects of physiology are assumed to be controlled by calcium in prokaryotes, including cell cycle progression, virulence, and competence (53–56). Calcium modulates the phosphorylation state of the Che proteins, which are the basis for chemotaxis (52). Our data suggest a link between chemotaxis, motility, and lanthanides.

Our nescience relating to the physiological role of calcium in prokaryotes includes calcium uptake. Complexes of short-chain polyhydroxybutyrate (PHB, a common polyhydroxyalkanoate [PHA]) and polyphosphate (PP) form calcium channels and represent one important route for Ca uptake (57–61). The sensitivity of these complexes to La was exploited to characterize them (62). Long chains of PHB, kept in a vacuole or inclusion body, are a common form of carbon storage. In previous work, we showed intracellular, periplasmic lanthanum accumulation in strain AL1 (32), often in close proximity to PHB vacuoles. We observed comparable periplasmic deposits when we grew AL1 with Nd or the Ln cocktail. We also noted differential expression of genes linked to PHB synthesis and PHB vacuole formation. An involvement of complexed PHB in selective lanthanide uptake (and storage) could explain the localization of the observed lanthanide deposits. Periplasmic calcium accumulation is a strategy for regulating intracellular calcium levels (63).

Differences in the composition of the observed Ln deposits suggest different mechanisms for accumulating different lanthanide elements. We did not detect P in periplasmic deposits from cultures grown with La. Determined P to Ln ratios for Nd and Ln cocktail samples align with potential lanthanide phosphates. For *M. extorquens* AM1, it was postulated that lanthanides are kept intracellularly in the cytoplasm complexed with polyphosphate (25). AL1 biomass samples originating from Ln cocktail cultures were previously shown to be enriched with heavier lanthanides (32), but this whole cell analysis did not allow to distinguish between lanthanides bound extracellularly and present within cells. The deposits from Ln cocktail cultures described here contain mostly Ce (38 Ln%), in nearly equal quantities (14-16 Ln%) Nd, Dy, Ho, and Er; but neither La and hardly any Yb. Periplasmic lanthanide deposition might function as a buffer. Together with a potential selective uptake into the cytoplasm, this could ensure that only preferably utilized lanthanides, for instance La, are incorporated into proteins.

The release of organic acids such as citric, oxalic, tartaric, and gluconic acid, forms the basis for heterotrophic bioleaching (64,65). Polyhydroxyl carboxylic acids such as gluconolactone can chelate metals, including lanthanides (66,67). One of the most strongly upregulated genes in response to Nd and the Ln cocktail encodes a FAD-dependent glucose 1-dehydrogenase (RHAL1_01212), which catalyzes the reaction β-D-glucose + NAD(P)^+^ ↔ gluconolactone + NAD(P)H + H^+^. Under non-methylotrophic growth conditions, the gene was downregulated when lanthanum was added (32). We have no robust data if gluconolactone is secreted and functions as a chelator in strain AL1. Our data warrant further investigations on organic acids being used by strain AL1 in the context of lanthanide chelation and uptake. If it functions as a chelator, the upregulation of the glucose dehydrogenase gene might indicate that Nd can only be taken up chelator-based. Chelating heavier lanthanides can reduce the uptake of non-utilizable lanthanides, for instance by blocking porins.

High intracellular concentrations of (non-)utilizable metals cause toxicity and must be avoided by homeostatic mechanisms. In the case of Nd and Ln cocktail incubations, simultaneous upregulation of heavy metal efflux mechanisms and down-regulation of Fe uptake systems suggests that lanthanides might be accidentally taken up by the latter. The simultaneous release of a chelator and downregulation of the machinery needed for the uptake of metal-chelator complexes was described as a strategy to deal with elevated copper levels in *P. aeruginosa (68)*. The downregulation of *ebbB* and *ebbD,* coding for part of the machinery needed for transmitting energy between cytoplasmic and outer membrane (69), in AL1 in response to Nd and the Ln cocktail indicates that the energy coupling needed to drive TonB-dependent transport is reduced. However, this seems inconsistent if Nd uptake is chelator-based.

We have no data supporting the involvement of lanthanides in sulfur metabolism, but it is noteworthy that *ssuABCD* are responsive to lanthanum concentration and different lanthanide elements. SsuABCD are responsible for taking up sulfonates and sulfonate esters. The differential expression of *ssuABCD* usually indicates sulfur starvation (70). SsuD was postulated as an indicator of oxidative stress under sulfur-limited conditions (71).

## CONCLUDING REMARKS

More and more pieces of the “lanthanide puzzle” (72) are identified. The lanthanome, the entirety of biomolecules, especially proteins, partaking in Ln-dependent metabolism (73), becomes more extensive. We showed that different lanthanide elements affect many genes, tied to various aspects of metabolism in Beijerinckiaceae bacterium RH AL1 and that strain AL1 can distinguish between different (utilizable) lanthanides. Not all of our findings do implicate causality, but they support the possibility that lanthanides can play a diverse role in bacterial physiology. Taking into account the relevance of lanthanides for our modern way of living, understanding these roles will help to tune lanthanide-dependent metabolism towards biotechnological applications.

## MATERIALS AND METHODS

### Cultivation

Beijerinckiaceae bacterium RH AL1 was grown using MM2 medium (74). Incubations were done at room temperature while shaking (110 rpm). Pre-cultures were grown on sodium pyruvate (0.2% wt/vol) as the carbon source in acid-washed 200 mL serum bottles sealed with boiled and sterilized butyl rubber stoppers. Biomass was collected by centrifugation and repeatedly washed before being used as inoculum. Cultivation experiments were performed in triplicates in acid-washed 150 mL Erlenmeyer flasks with cellulose stoppers and methanol (0.5 % vol/vol) as the carbon source. Cultures were supplemented with either different concentrations of lanthanum (50 nM, 1 µM) or different lanthanide elements (La, Nd, lanthanide cocktail [La, Ce, Nd, Dy, Ho, Er, Yb]; 1 µM). Cultures were grown to the stationary phase, and samples for downstream, RNAseq-based transcriptome analysis were taken during the mid-exponential phase. Additional samples were taken for cell counts based on quantitative PCR (qPCR).

#### DNA extraction and quantitative PCR (qPCR)

Genomic DNA was extracted following standard protocols using the NucleoSpin Microbial DNA Mini kit (Macherey-Nagel, Düren, Germany). DNA was quantified using a Qubit 3 fluorometer in combination with dsDNA BR or HS assay kits (ThermoFisher Scientific, Frankfurt, Germany). DNA integrity was checked through spectrophotometry and DNA agarose gel electrophoresis. Details concerning qPCR can be found in the **supplementary material**.

#### RNA extraction, mRNA enrichment, and sequencing library preparation

Total RNA was extracted and mRNA enriched from biomass samples collected in the middle exponential growth phase based on previously described methods (32,75). Sequencing libraries were prepared with the NEBNext Ultra II Directional RNA library prep kit for Illumina (New England Biolabs, Ipswich, Massachusetts, USA) and checked by chip-based, high-resolution gel electrophoresis with a Bioanalyzer instrument and DNA 7500 Pico reagents (Agilent Technologies, Santa Clara, California, USA).

#### Sequencing and data pre-processing

An equimolar pool of libraries was subjected to Illumina sequencing (2 ✕ 100 bp, paired ends) with a NovaSeq 6000 instrument and an SP flowcell (Illumina, San Diego, California, USA). Sequencing was carried out by the sequencing core facility of the Leibniz Institute on Aging - Fritz Lipmann Institute. The quality of raw and later trimmed sequences was checked with *FastQC* (v0.11.9) (76). Data pre-processing was done as described previously (32) and is described in the **supplementary material**.

#### Differential gene expression analysis

Differential gene expression analysis was done using the packages *edgeR* (v3.20.9) (77), *limma* (v3.50.0) (78), *mixOmics* (v6.18.1) (79), *HTSFilter* (v1.34.0) (80), and *bigPint* (v1.10.0) (81), in the R software framework for statistical computing (v3.5.1) (82). Further details are given in the **supplementary material.** Differentially expressed genes were filtered concerning the log_2_ fold change (log_2_FC), FDR-corrected *P* value (**Figure S4**), and absolute gene expression in log_2_counts per million (cpm). Genes with changes in gene expression above |0.58| log_2_FC (fold change), gene expression values higher than 4 (log_2_CPM, counts per million), and p-values smaller than 0.05 were considered for downstream analysis.

#### Overrepresentation analysis, gene set enrichment analysis, analysis of unknown and hypothetical proteins

Subsets of differentially expressed genes were subjected to overrepresentation analysis and GSEA using the functions *enrichKEGG* (*p*-value cutoff 0.05, *p*-value adjustment Bejamini-Hochberg procedure) and *gseKEGG* (*p-*value cutoff 0.05, *p*−value adjustment Bejamini-Hochberg procedure) from the R package *clusterProfiler* (v. 4.4.3) (83,84) and the available genome annotation from the KEGG GENOMES databases (identifiiers: T06029, “bbar”). The translated amino acid sequences of genes coding for hypothetical and proteins of unknown function were queried against NCBI nr (85), Pfam (86), and Interpro (87) applying the default settings of available web services.

#### Motility assay

To investigate the motility of Beijerinckiaceae bacterium strain RH AL1 when grown with different lanthanide elements and concentrations (La, Nd, Ln cocktail; 5 nM, 10 nM, 50 nM, 100 nM, 1 µM, 5 µM, 10 µM). Incubations were done with MM2 medium (74), semi-solidified with agar (0.4%, w/v), and containing 2,3,5-triphenyltetrazolium chloride (0.005 %, w/v) (Carl Roth GmbH & Co. Kg, Karlsruhe, Germany) (88). Incubations were done in 6-well plates with methanol (0.5%) as carbon and energy source. A preculture grown to stationary phase with pyruvate (0.2%, w/v) as a carbon source and in absence of lanthanides was washed five times with 10 mL basal MM2 medium and used as an inoculation source for the motility assay. The evaluation was performed 14 days after inoculation.

#### Electron microscopy

Sample preparation for transmission electron microscopy (TEM), freeze-fracture TEM, and Energy-dispersive X-ray spectroscopy (EDX) analyses were carried out according to (32). Sample preparation is outlined in the **supplementary material**. The deconvolution of EDX spectra was done using the Quantax software (Bruker, Berlin, Germany).

#### Analysis of TEM pictures

Cell areas and the areas occupied by PHB vacuoles were determined using *ImageJ* (v. 1.52r) (89,90) and its freehand selection tool. The analysis was done for three images (magnification: 4000 ✕, image area: 540.5 µm^2^), and between 27 to 30 cells were analyzed per condition (La, Nd, cocktail).

#### Statistical analysis

Motility-assay data were tested for normal distribution (*Kolmogorov-Smirnov test*), outliers (*Dean-Dixon test*) and trend (*Neumann trend test*). Statistical analysis was performed with Microsoft Excel based on a 95% confidence Interval.

#### Figure generation

Plotting was done with the R software framework (v. 4.2.1) (91) using the packages *ggplot2* (v. 3.3.6) (92), *gplots* (v. 3.1.3) (93), and *cowplot* (v. 1.1.1) (94), and *upsetR* (v. 1.4.0) (95), including their respective dependencies. Figures were finalized with inkscape (https://inkscape.org/).

#### Data availability

RNA-seq data sets can be accessed via EBI/ENA ArrayExpress (accession: E-MTAB-12015 [https://www.ebi.ac.uk/arrayexpress/experiments/E-MTAB_12015/]). The genome of Beijerinckiaceae bacterium RH AL1 is available via the EBI/ENA accession numbers: LR590083 (https://www.ebi.ac.uk/ena/data/view/LR590083) and LR699074 (https://www.ebi.ac.uk/ena/data/view/LR699074) (genome and plasmid). We also provide details about sequence data processing and differential gene expression analysis via the Open Science Framework (https://osf.io/) (https://osf.io/p2nf6/?view_only=b83c7bbd806b43bdac419ebc8117eaa0).

## ACKNOWLEDGEMENTS

CEW thanks N. Cecilia Martinez-Gomez and Nathan M. Good (UC Berkeley, CA, USA) for sharing and discussing unpublished data. This research was supported by the Deutsche Forschungsgemeinschaft (grant: WE6579/4-1, granted to CEW).

## AUTHOR CONTRIBUTIONS

LG designed, and carried out experimental work, performed RNAseq data analysis, and wrote a first draft of the manuscript. JB helped with experimental work. MW and FS did all the EM work and helped with interpreting EM data. CEW designed the experimental work, carried out RNAseq data pre-processing and data analysis, acquired funding, and wrote the manuscript based on input from all co-authors.

