## Supplementary Material for "Different lanthanide elements induce strong gene expression changes in a lanthanide-accumulating methylotroph"

***SUPPLEMENTARY MATERIAL:*** **Different lanthanide elements induce strong gene expression changes in a lanthanide-depositing methylotroph**

Linda Gorniak[1], Julia Bechwar[1], Martin Westermann[2], Frank Steiniger[2], Carl-Eric Wegner[1,#]

[1]Institute of Biodiversity, Aquatic Geomicrobiology, Friedrich Schiller University, Dornburger Str. 159, 07743 Jena, Germany

[2]Electron Microscopy Center, Jena University Hospital, Ziegelmühlenweg 1, 07743 Jena, Germany

### Corresponding author:

Carl-Eric Wegner

Running title: Lanthanides cause strong gene expression changes

Keywords: lanthanides, lanthanome, RNAseq, EDX, TEM, FFTEM

**SUPPLEMENTARY MATERIAL**

**Supplementary Information.** Complementary information about used materials and applied methods.

We also provide details about sequence data processing and differential gene expression analysis via the Open Science Framework (https://osf.io/) (https://osf.io/p2nf6/?view_only=b83c7bbd806b43bdac419ebc8117eaa0)

*Quantitative PCR*

DNA was diluted, and one to 20 ng of genomic DNA was used as a template for qPCR. The copy numbers of the *lanM* gene (coding for lanmodulin) were determined using a CFX96 instrument (Bio-Rad, Munich, Germany), Brilliant II SYBR® Green QPCR Master Mix (Agilent Technologies, Germany) to deduce cell numbers. *lanM* is a single-copy gene in Beijerinckiaceae bacterium RH AL1. We designed a primer set *lanM_f1/lanM_r1* (*lanM_f1:* 5’-GGATTTTCGCAAGACCCTTT-3’, *lanM_r1:* 5’-CTTCTTGAACTTGGCCTTGG-3’) and used it with the following cycling conditions: 15 min. denaturation at 94°C, followed by 30 cycles of 30 s at 94°C, 60 s at 55°C, and 45 s at 72°C. Plasmids containing a *lanM* fragment from strain AL1 were used for standard curves. Standard curves were linear from $5 \times 10^8$ to $5 \times 10^2$ copies with $R^2$ values and PCR efficiencies above 0.99 and 80%, respectively. Dilution series were used to assess potentially present PCR inhibitors. Melt curve analysis was done to verify PCR specificity. qPCR was done based on three biological replicates and three technical replicates per biological replicate.

*RNAseq data pre-processing*

Adaptor- and quality-trimming (settings: minlen= 75, qtrim = rl, ktrim = rl, k = 25, mink= 11, trimq = 20, qtrim = rl) were carried out with *bbduk* (v38.26) (1) using its included database of common sequence contaminants and adapters. Trimmed reads were filtered with SortMeRNA (v2.1) (2) and the SILVA (3) and Rfam (4) databases by removing rRNA-derived and non-coding RNA sequences. Read mapping onto the available reference genome of Beijerinckiaceae bacterium RH AL1 (EBI accession no. LR590083 [genome] and LR699074 [plasmid]) (5) was done with *bbmap* (v.38.26) (settings: slow, k = 11). The number of mapped reads per feature (e.g. coding genes) was determined from *.bam* files, that have been sorted and indexed with *samtools* (v1.3.1) (6), using *featureCounts,* which is part of the *Subread* package (v1.6.3) (7, 8).

*Differential gene expression analysis*

Pseudo-counts ($\log_2$(counts+1)) (**Figure S1**) were generated for subsequent data exploration by means of MA (mean of the normalized counts versus the $\log_2$FC [fold change] for all genes tested) plots. Multidimensional scaling plots were generated using the *plotMDS* function of *limma* (v. 3.50.0) (9). Inter-dataset relationships were in addition assessed based on the hierarchical clustering of scaled read count data (**Figure S2**). Scatterplot matrices were plotted with the *plotSM* function of *bigPint* (v. 1.10.0) (10) to check how replicates of conditions behaved when compared to each other and to the replicates of other conditions. Genes with insufficient read count data were filtered with the *filterByExpr* function which is part of *edgeR* (v. 3.20.9) (11). Libraries were normalized by calculating normalization factors and estimating dispersion. The biological coefficient of variation (BCV) (**Figure S3**) was used to identify the uncertainty regarding transcript abundance based on replicate groups. P-values derived from the exact test proposed by Robinson and Smyth (13) have been assessed for all carried out comparisons (**Figure S4**)

*Transmission elektron microscopy (TEM)*

Biomass was harvested by centrifugation and fixed overnight at 4°C by use of glutaraldehyde (2.5 %, v/v) in cacodylate buffer (100 mM, pH 7.3). Cells were washed with cacodylate buffer three times and incubated 5, 10, and 15 minutes prior to pelleting the biomass. All centrifugation steps were performed at room temperature and 4000 × g for 10 minutes. The pellets were resuspended and kept in 500 ml cacodylate buffer at 4°C. Further processing was according to Wegner et al., 2021 (12). Fixed samples were dehydrated in an ethanol series and stained with 2% (w/v) uranyl acetate in 50% (v/v) ethanol. Araldite resin (Plano, Wetzlar, Germany) was used for embedding samples. Ultrathin sections (70 nm thickness) were cut using an ultramicrotome Ultracut E (Reichert-Jung, Vienna, Austria) and mounted on Formvar-carbon coated 100 mesh grids (Quantifoil, Großlöbichau, Germany). The ultrathin sections were stained with lead nitrate for 10 minutes, examined in a Zeiss CEM 902 A electron microscope (Carl Zeiss AG,Oberkochen, Germany), and imaged using a TVIPS 1k Fast-Scan CCD-Camera (TVIPS, Munich, Germany). Ultrathin sections and freeze-fracture replicas were examined in a digital Zeiss EM 900 electron microscope (Zeiss, Oberkochen, Germany; digital upgrade by Point Electronic, Halle, Germany) operated at 80 kV. Digitized images were taken with a wide-angle dual-speed 2K CCD camera controlled by the Sharp:Eye base controller and operated by the Image SP software (TRS, Moorenweis, Germany).

*Freeze fracture transmission electron microscopy (FFTEM)*

Glutaraldehyde-fixed biomass was supplied with the cryoprotectant glycerol (final concentration 15%), and 2 µl samples were transferred into a pair of copper hat-type carriers. Copper sandwiches were incubated for 1 minute in a cooled propane-ethane mix to rapidly plunge-freeze the samples before transferring them into liquid nitrogen. A BAF400T freeze fracture unit (BAL-TEC, Liechtenstein) supplied with a double-replica stage was used to freeze-fracture the samples at -110°C and high vacuum conditions ($<10^{-4}$ Pa). Platinum/carbon replicates were created by use of an electron beam vaporizer (1350 V - 1950 V, 89 mA - 93 mA), by adding a 2 nm platin layer at a 35° angle onto the fractured surface, followed by vertical evaporation with coal (ca. 20 nm layer). Cell material was removed from replicas by use of sodium dodecyl sulfate. Replicas were washed four times in distilled water and transferred onto uncoated EM-grids for examination in a TEM as outlined above.

**Fig S1.** Comparison of pseudocount distributions between RNAseq data sets. Pseudocounts were determined gene-wise by calculating $Log_2$([count of mapped reads]+1).

**Fig S2.** Hierarchical clustering of RNAseq data sets. The clustering was based on a distance matrix that was generated by subtracting the Spearman correlation between data sets from 1. The resulting distances ranged between 0 and 2. Spearman correlations were calculated from scaled cpm (counts per million).

**Fig S3.** Biological coefficient of variation (BCV) plot. The BCV indicates the coefficient of variation with which the true abundance of the gene transcripts varies between replicate RNAseq data sets. $Log_2CPM = log_2$ counts per million

**Fig S4.** P-value distributions resulting from testing for differential gene expression between groups of RNAseq data sets. The p-values were derived from the exact test proposed by Robinson and Smyth (13) and implemented in *edgeR (11)*.

**Fig S5.** Soft-agar based motility assay. Fourteen days after inoculation, diameters of cell intrusion into the soft-agar medium were measured and respective areas were calculated. For each condition (supplementation with La, Nd, or lanthanide cocktail), increasing lanthanide concentration led to a decrease in motility.

**Fig S6.** Deconvolution of EDX spectra. The deconvolution for lanthanum deposits was done based on measurements published previously (12) (left panel), and the right panel shows the deconvolution for an exemplary Nd deposit (A). The deconvolution for the deposit originating from cells grown with the $Ln^{3+}$ cocktail is shown for two different energy ranges. The panel on the right provides a more zoomed-in representation of the spectrum. We did three replicate measurements for each condition using different deposits and cells (**Table S12**).

**Table S1.** Quantitative PCR targeting the 16S rRNA gene and lanM.

**Table S2.** Overview of gene expression data of Beijerinckiaceae bacterium RH AL1 grown with methanol (0.5%, v/v) as carbon source, and supplemented with either 50 nM or 1 µM La, 1 µM Nd, or an equimolarly pooled (1µM) cocktail of light and heavy lanthanides (La, Ce, Nd, Dy, Ho, Er, Yb). Gene expression is given in log2 counts per million (CPM). La.1-3 = RH AL1 grown with 1 µM La, Nd.1-3 = RH AL1 grown with 1 mM Nd, Cocktail.1-3 = RH AL1 grown with an equimolarly pooled cocktail of heavy and light lanthanides (1 µM), La_low.1-3 = RH AL1 grown with 50 nM La, La_high.1-3 = RH AL1 grown with 1 µM. The latter two groups of data sets originate from a different incubation than the first free groups of data sets.

**Table S3.** Overview of differentially expressed genes in Beijerinckiaceae bacterium RH AL1 when grown with either 50 nM or 1 µM La. logFC = $log_2$ Foldchange, logCPM = $log_2$ Counts per million, FDR = false discovery rate.

**Table S4.** Overview of differentially expressed genes in Beijerinckiaceae bacterium RH AL1 when grown with either 1 µM La or 1 µM Nd. logFC = $log_2$ Foldchange, logCPM = $log_2$ Counts per million, FDR = false discovery rate.

**Table S5.** Overview of differentially expressed genes in Beijerinckiaceae bacterium RH AL1 when grown with either 1 µM La or 1 µM of an equimolarly pooled lanthanide cocktail (La, Ce, Nd, Dy, Ho, Er, Yb). logFC = $\log_2$ Foldchange, logCPM = $\log_2$ Counts per million, FDR = false discovery rate.

**Table S6.** Overview of differentially expressed genes in Beijerinckiaceae bacterium RH AL1 when grown with either 1 µM Nd or 1 µM of an equimolarly pooled lanthanide cocktail (La, Ce, Nd, Dy, Ho, Er, Yb). logFC = $\log_2$ Foldchange, logCPM = $\log_2$ Counts per million, FDR = false discovery rate.

**Table S7.** Overrepresentation and gene set enrichment analysis. Based on KEGG annotations and gene sets shared or unique for individual comparisons, we determined overrepresented/enriched functions with *clusterProfiler* (14, 15).

**Table S8.** Presence-absence matrix for differential gene expression in response to different lanthanide elements and lanthanum concentration. To summarize the results, only genes associated with selected aspects of metabolism were included, e.g. motility and chemotaxis, carbohydrate metabolism, PHA cycle, and transport.

**Table S9.** Results from the carried out motility assay with diameters and calculated areas of cell intrusion into the CMM2 soft-agar medium after 14 days of incubation. Data series (La, Nd, lanthanide cocktail) were normally distributed (p-value < 0.29, *Kolmogorov-Smirnov test*) without outliers (p-values < 0.29, *Dean-Dixon test*) and showed a decreasing trend (p-value < 0.89, *Neumann trend test*) of motility with increasing lanthanide concentrations. Statistic tests were performed based on a 95% confidence interval.

**Table S10.** Overview of gene expression data for lanthanome(-related genes). Gene expression is given in log2 counts per million (CPM). Gene name refers to the gene name in M. extorquens AM1. Genes were grouped as described in the main text. Gene annotation data was taken from Wegner et al. 2020 (5).

**Table S11.** Lanthanome(-related) genes in *M. extorquens* AM1 and Beijerinckiaceae bacterium RH AL1. Gene homologs in strain AL1 have been identified by *blastp* (16) searches of AM1 amino acid sequences against a protein database containing all amino acid sequences encoded in the genome of strain AL1. The queried sequences comprised relevant gene products identified previously (17), especially gene products of the lut-cluster, as well as LC cluster gene products described by Zytnick and colleagues (18). pident = percentage identity, qstart = query start, qend = query end, sstart = subject start, send = subject end.

**Table S12.** Results from the deconvolution analysis of obtained EDX spectra. EDX-based elemental analysis was done for periplasmic deposits identified in cells grown with Nd (n=3), the lanthanide cocktail (n=1), and La (n=3, Wegner et al., 2021). At % = atomic percent, Ln % = proportion of the respective Ln based on the sum of the At % for all Ln.

**<u>Note:</u>** All supplementary tables are provided in one combined spreadsheet.

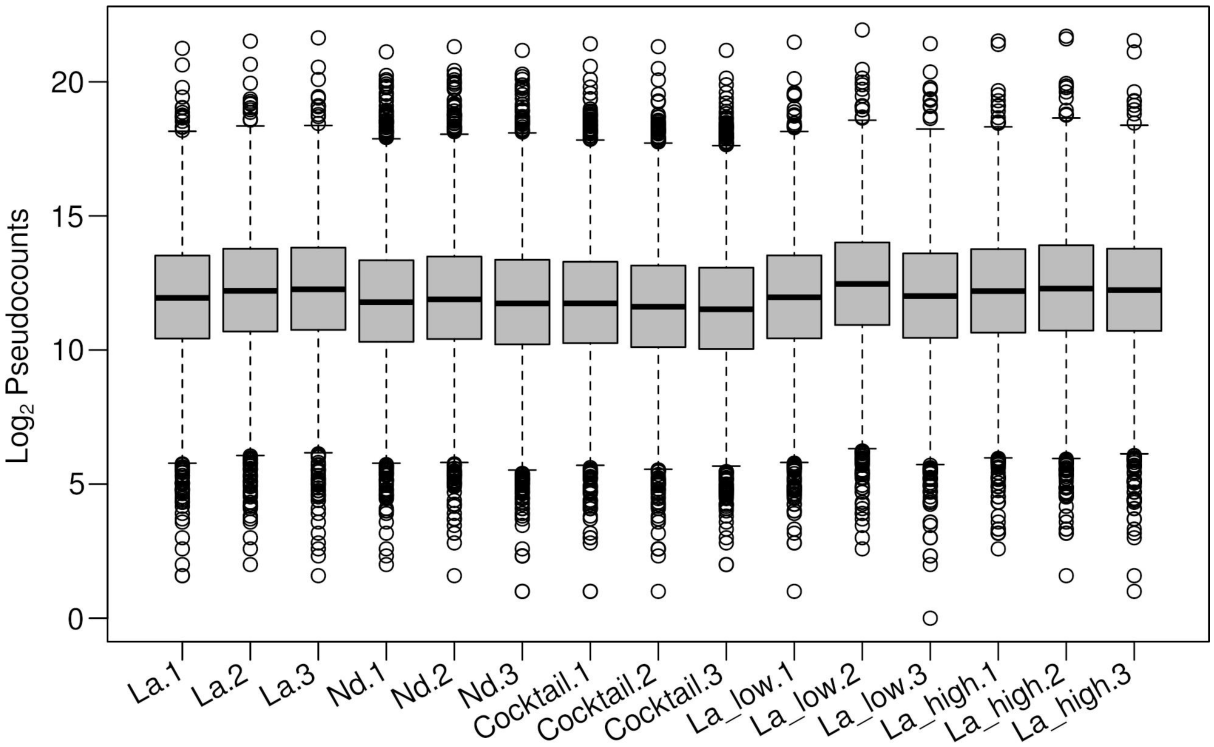
Log2 Pseudocounts
0
5
10
15
20
La.1
La.2
La.3
Nd.1
Nd.2
Nd.3
Cocktail.1
Cocktail.2
Cocktail.3
La_low.1
La_low.2
La_low.3
La_high.1
La_high.2
La_high.3

**Fig. S1**

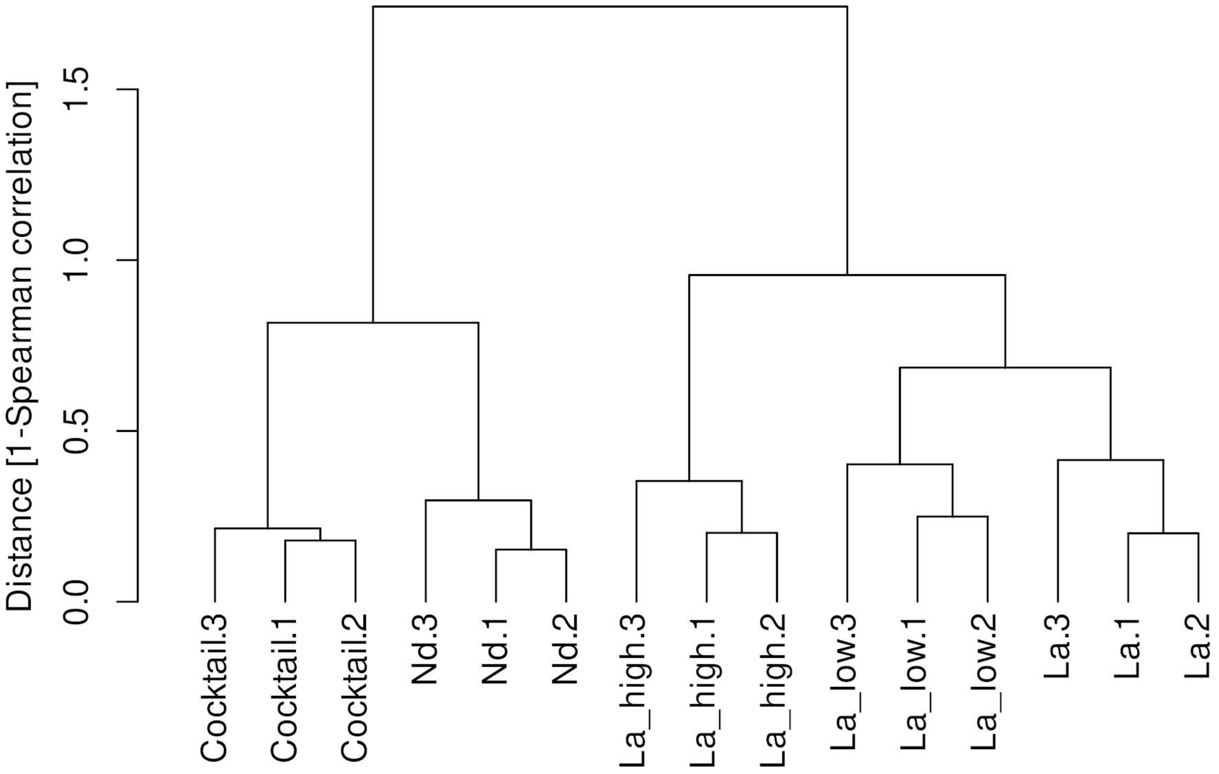

Distance [1-Spearman correlation]
0.0
0.5
1.0
1.5
Cocktail.3
Cocktail.1
Cocktail.2
Nd.3
Nd.1
Nd.2
La_high.3
La_high.1
La_high.2
La_low.3
La_low.1
La_low.2
La.3
La.1
La.2


**Fig. S2**

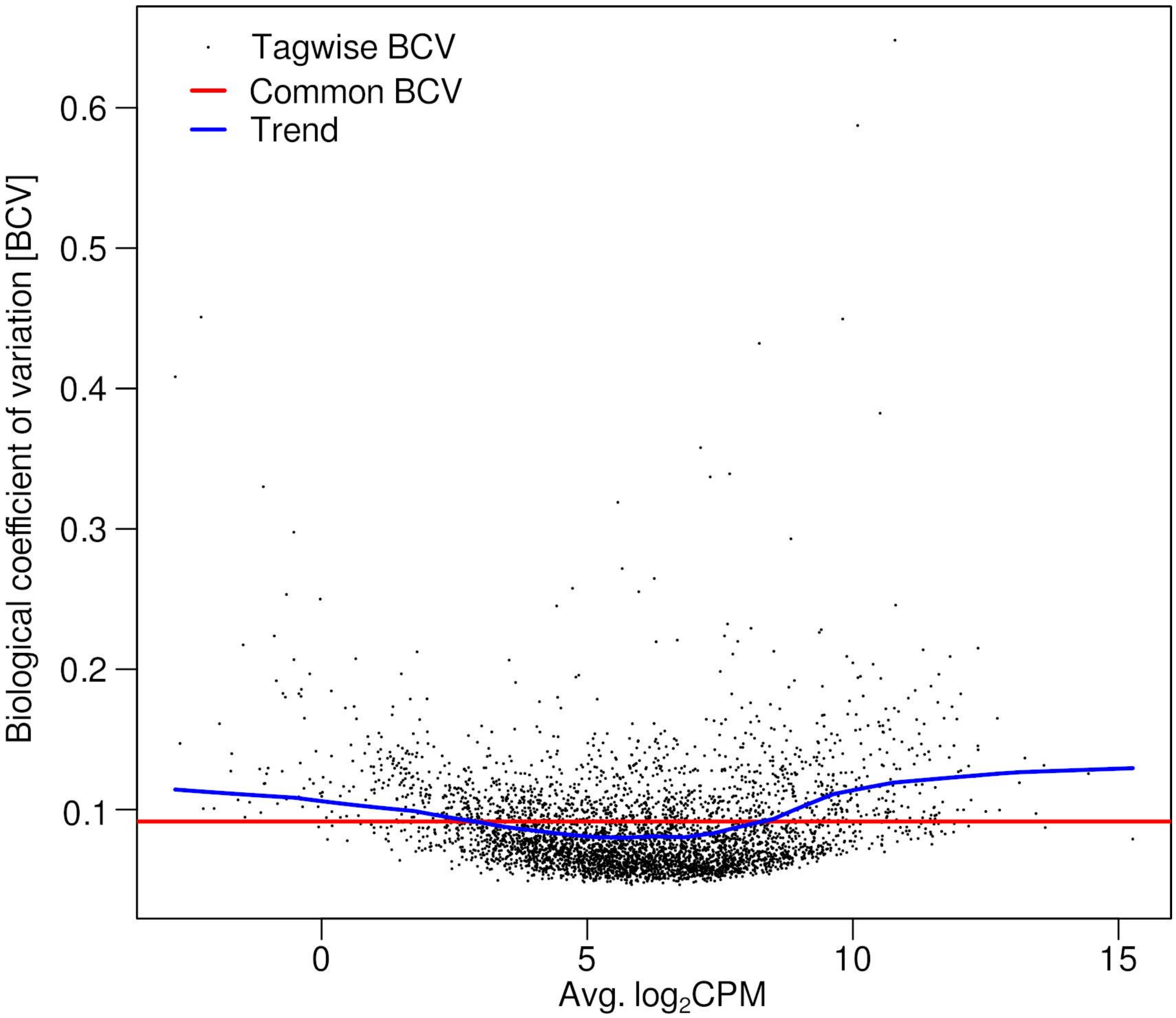

Tagwise BCV
Common BCV
Trend
0.6
0.5
0.4
0.3
0.2
0.1
Biological coefficient of variation [BCV]
0
5
10
15
Avg. log2CPM


**Fig. S3**

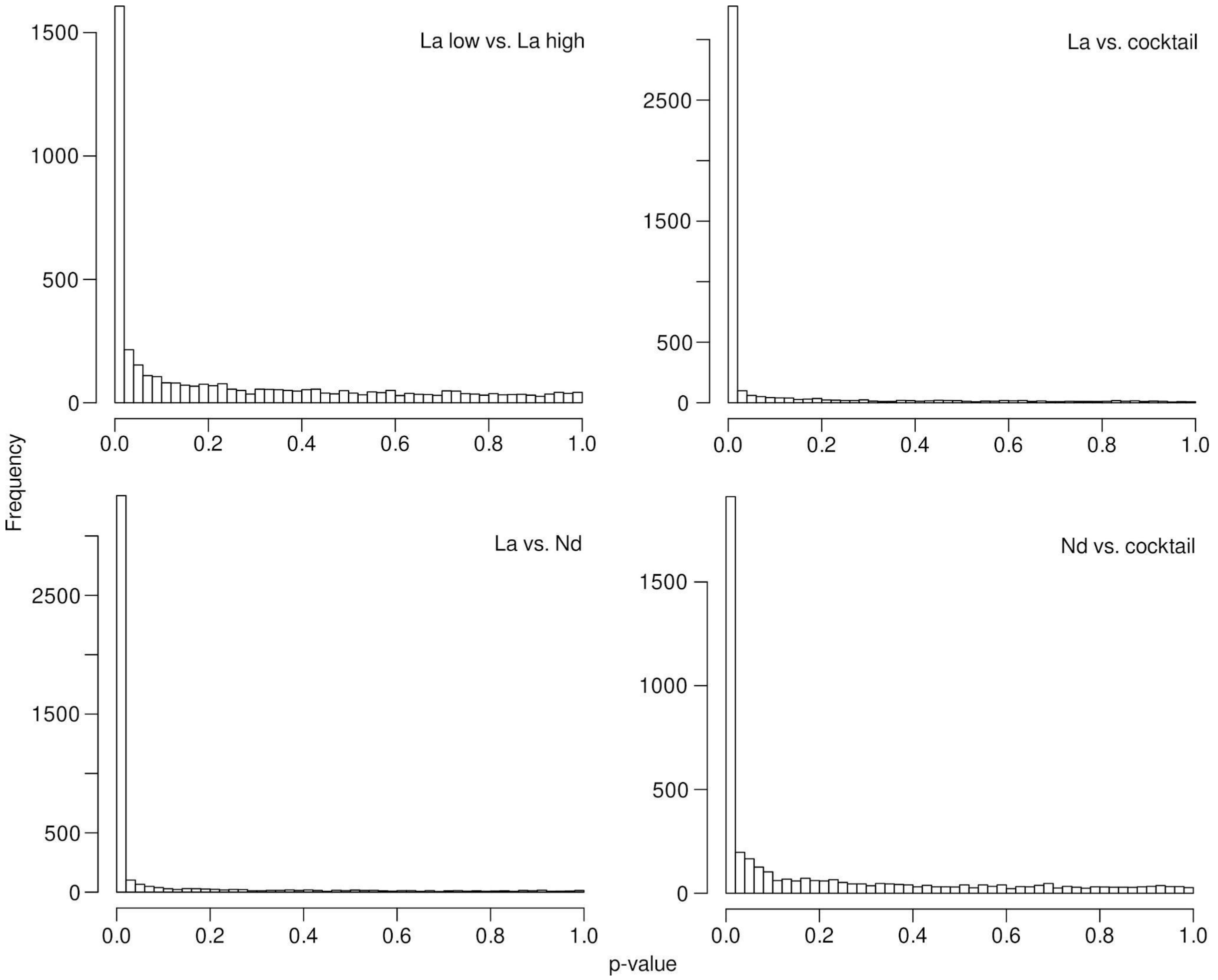

La low vs. La high
La vs. cocktail
La vs. Nd
Nd vs. cocktail
Frequency
p-value
0
500
1000
1500
2500
0.0
0.2
0.4
0.6
0.8
1.0


**Fig. S4**

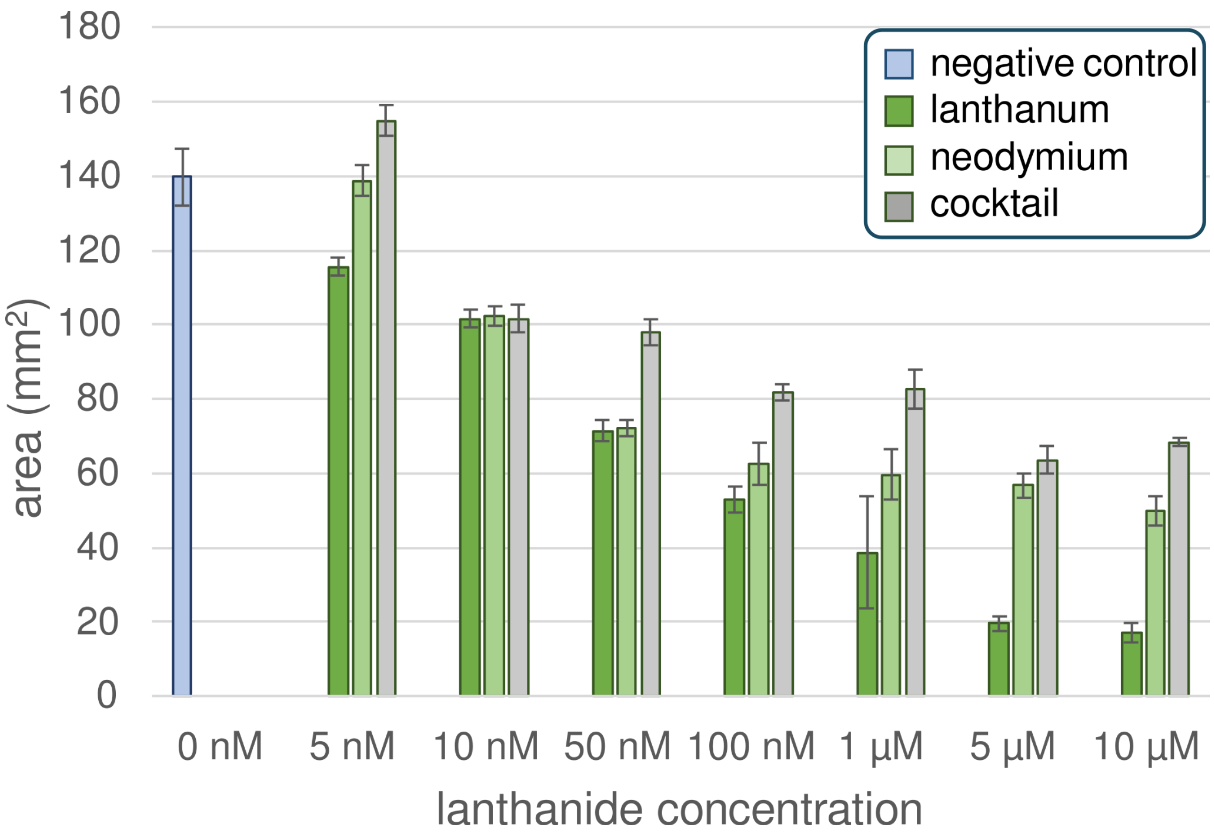
negative control
lanthanum
neodymium
cocktail
180
160
140
120
100
80
60
40
20
0
area (mm²)
0 nM
5 nM
10 nM
50 nM
100 nM
1 µM
5 µM
10 µM
lanthanide concentration

**Fig. S5**

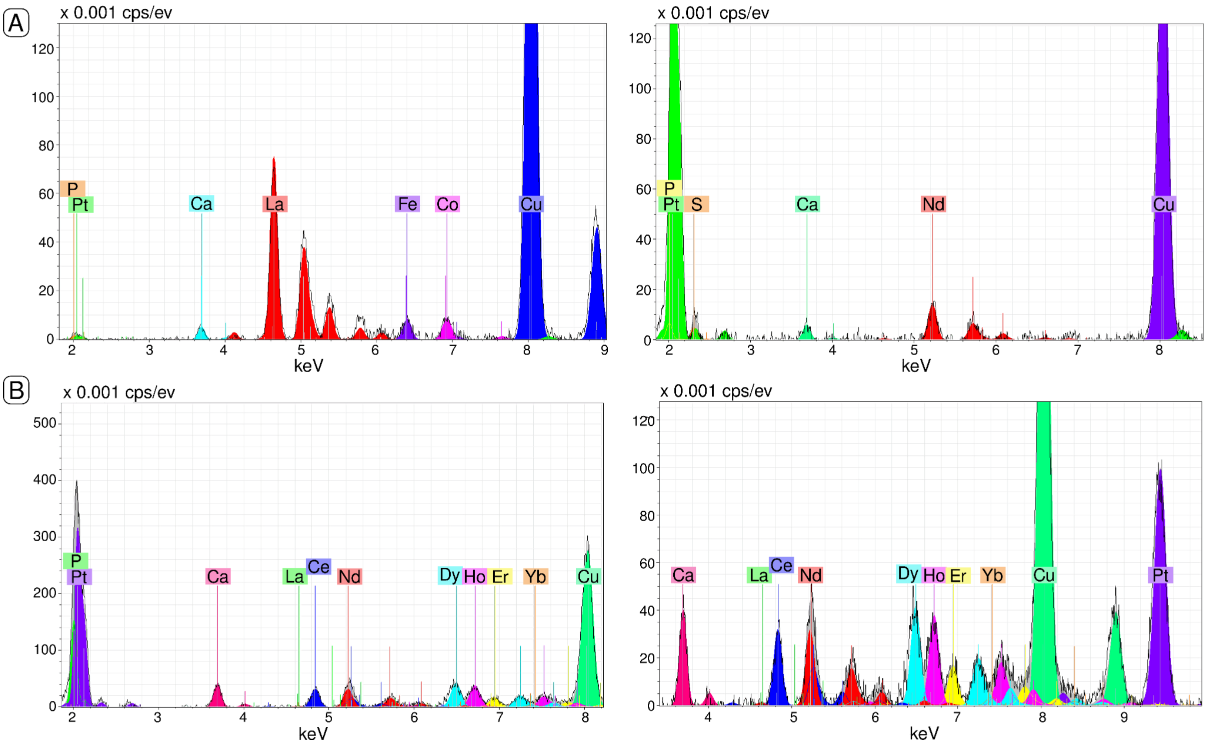
A
x 0.001 cps/ev
P
Pt
Ca
La
Fe
Co
Cu
keV
x 0.001 cps/ev
P
Pt
S
Ca
Nd
Cu
keV
B
x 0.001 cps/ev
P
Pt
Ca
La
Ce
Nd
Dy
Ho
Er
Yb
Cu
keV
x 0.001 cps/ev
Ca
La
Ce
Nd
Dy
Ho
Er
Yb
Cu
Pt
keV

**Fig. S6**
